# Decoding nucleosome positions with ATAC-seq data at single-cell level

**DOI:** 10.1101/2021.02.07.430096

**Authors:** Bingxiang Xu, Xiaoli Li, Xiaomeng Gao, Yan Jia, Feifei Li, Zhihua Zhang

## Abstract

As the basal bricks, the dynamics and arrangement of nucleosomes orchestrate the higher architecture of chromatin in a fundamental way, thereby affecting almost all nuclear biology processes. Thanks to its rather simple protocol, ATAC-seq has been rapidly adopted as a major tool for chromatin-accessible profiling at both bulk and single-cell level. However, to picture the arrangement of nucleosomes *per se* remains a challenge with ATAC-seq. In the present work, we introduce a novel ATAC-seq analysis toolkit, named deNOPA, to predict nucleosome positions. Assessments showed that deNOPA not only outperformed state-of-the-art tools, but it is the only tool able to predict nucleosome position precisely with ultrasparse ATAC-seq data. The remarkable performance of deNOPA was fueled by the reads from short fragments, which compose nearly half of sequenced reads and are normally discarded from nucleosome position detection. However, we found that the short fragment reads enrich information on nucleosome positions and that the linker regions were predicted by reads from both short and long fragments using Gaussian smoothing. We applied deNOPA to a single-cell ATAC-seq dataset and deciphered the intrapopulation heterogeneity of the human erythroleukemic cell line (K562). Last, using deNOPA, we showed that the dynamics of nucleosome organization may not directly couple with chromatin accessibility in the cis-regulatory regions when human cells respond to heat shock stimulation. Our deNOPA provides a powerful tool with which to analyze the dynamics of chromatin at nucleosome position level in the single-cell ATAC-seq age.

## Introduction

Eukaryotic genomes are packaged into chromatin, which consists of repeating nucleosomes that consist of a histone octamer wrapped around 147 bp of DNA and serves as the fundamental building blocks for higher chromatin architecture.

The *in vivo* nucleosome positions are highly dynamic, being influenced by, for example, chromatin remodelers (Rippe et al. 2007), the binding of transcription factors (Nie et al. 2014) and transcription (Valouev et al. 2011). Thus, the positioning and occupancy of nucleosomes contribute to the heterogeneity and flexibility of chromatin between cell types, as well as among cell populations. However, the positioning and occupancy of nucleosomes are not completely randomly distributed in the genome. In addition to DNA sequence preferences (Kaplan et al. 2009), well-defined patterns are found in cis-regulatory elements, e.g., the nucleosome free region (NFR), the well-phased flanking nucleosome arrays in promoters (Ozsolak et al. 2007; Mavrich et al. 2008a; Mavrich et al. 2008b), and the bimodal at some enhancers (He et al. 2010). Periodic arrangement patterns are also found at the binding loci of transcription factors, e.g., CTCF (Fu et al. 2008), characterizing the protein and chromatin context (Zhang et al. 2017). On the other hand, the nucleosomes may also modulate transcription factor (TF) binding and transcription machinery by their positioning (Lickwar et al. 2012). Therefore, nucleosome arrangement is tightly linked with gene regulation (Schones et al. 2008) and the response to various endogenous or exogenous stimulation, e.g., heat shock (Shivaswamy and Iyer 2008).

Experimental technologies have been developed to generate genome-wide nucleosome positions using high-throughput sequencing. For example, the chemical mapping approach captures nucleosome positions by sequencing DNAs that bind to genetically modified histone H4 (Brogaard et al. 2012). Since only DNA sequences binding to histone octamers were specifically read out (Brogaard et al. 2012), chemically mapped nucleosome position has been regarded as the gold standard (Brogaard et al. 2012; Voong et al. 2016). Micrococcal nuclease digestion with deep sequencing (MNase-seq) is another widely used technology, which utilizes micrococcal nuclease to cut the linker DNA between two neighboring nucleosomes (Pajoro et al. 2018). Assay for Transposase-Accessible Chromatin using sequencing (ATAC-seq), which utilizes Tn5 transposase to digest accessible genomic DNA, has now become widely adopted for chromatin accessibility detection and nucleosome positioning (Buenrostro et al. 2013). ATAC-seq has continued to attract attention in recent years by the simplicity of its operation and its high-quality data (Buenrostro et al. 2013; Corces et al. 2017). Most recently, the development of single-cell ATAC-seq technology has advanced and has become commercially available (Cusanovich et al. 2015), opening a window to understand the dynamics of chromatin at the single-cell level, e.g., the dynamics of chromatin during embryonic development (Cusanovich et al. 2018) and the evolution of hematopoiesis and leukemia (Corces et al. 2016).

A long list of computational tools for nucleosome positions or chromatin accessibility analysis using data from these experiments is available in the literature. These tools can largely be divided into two categories. The first category, including such tools as nucleoATAC (Schep et al. 2015), TemplateFilter (Weiner et al. 2010), NucleoFinder (Becker et al. 2013) and NucID (Zhong et al. 2016), capture the consensus signal profile from a series of predefined positive nucleosome sites (template). This template is then used to scan the genome to find sites with signal matches. It is worth noting that nucleoATAC is the only ATAC-seq data-based tool designed and trained in a high-quality dataset in *yeast* (Schep et al. 2015). The second category, including such tools as NPS (Zhang et al. 2008), iNPS (Chen et al. 2014), DANPOS (Chen et al. 2013), nucleR (Flores and Orozco 2011) and many more (Teif 2016), is template-free and detects nucleosomes by capturing the local geometry characteristics of MNase-seq data. With the growing interest in gene regulation studies using ATAC-seq assay (Duren et al. 2017; Miraldi et al. 2019), the demands for nucleosome position detection from such data have exploded. However, it remains challenging to demand nucleosome position with low-resolution ATAC-seq data.

To address this problem, we herein presented a scheme for **<u>de</u>**coding **<u>n</u>**ucleosome **<u>o</u>**rganization **<u>p</u>**rofile based on **A**TAC-seq data (deNOPA). We showed evidence that deNOPA can detect nucleosome positions with ultralow resolution ATAC-seq data. This extraordinary sensitivity was achieved by making full use of all sequencing reads, while most state-of-the-art tools utilize only part of the reads from relatively long fragments (Buenrostro et al. 2013; Rowley et al. 2017). We showed that reads from short fragments also carry rich information for nucleosome position prediction. The deNOPA was assessed by both gold standard chemical mapping data and high-resolution MNase-seq data. Further, we showed that the ultra-sensitivity of deNOPA can help to decipher the intra-population heterogeneity of cancer cells with single-cell ATAC-seq data. Finally, we applied deNOPA to a classical model system of gene regulation, i.e., the heat shock response of human cells, and found that heat shock strongly influences nucleosome organization, but leaves nucleosome free regions and their accessibility largely intact. These results imply that profiling nucleosome positions with deNOPA could be a powerful approach to decipher dynamics of chromatin under ultralow resolution ATAC-seq conditions.

## Results

### A novel ATAC-seq-based nucleosome positioning prediction algorithm termed deNOPA

We developed deNOPA to predict nucleosome positions using ultralow coverage ATAC-seq data. In addition to long-fragment ATAC-seq reads, deNOPA also utilizes short-fragment reads (e.g., <~ 100bp) for prediction. Although short-fragment reads normally consist of 40% to 50% of total reads in ATAC-seq libraries, they have been discarded in most state-of-the-art nucleosome positioning studies (Buenrostro et al. 2013; Schep et al. 2015), as only long-fragment reads were considered as having originated from cutting events at the nucleosome linkers (Buenrostro et al. 2013). However, we found that the Tn5 cutting sites, as defined by short and long ATAC-seq fragments, are highly correlated. The distribution of Tn5 cutting sites, i.e., the 5’ ends of reads along the genomes, was found to be highly correlated between short and long fragments in *yeast* (Schep et al. 2015) and human (Buenrostro et al. 2013) (R=0.7408 and 0.7202, both with P < 10^-52^, respectively, Supplemental Information (SI) and Fig. S1). Further, the cutting sites of both long and short fragments were found to be highly enriched around the chemically mapped nucleosome linkers (Brogaard et al. 2012) (SI and Fig. S1). Therefore, we reasoned that the retrieval of these short-fragment reads could substantially increase the effective total reads coverage without the requirement of additional sequencing.

Briefly, deNOPA workflow consists of four distinct steps (Fig. 1). First, deNOPA summarizes input data into the distribution of reads coverage and Tn5 cutting events. Second, the ATAC-seq reads enriched regions (denoted as ARER) were called. Third, the candidate nucleosome positions were determined as loci with two flanking linkers having genomic distance between 101 to 215bp. The distance threshold we used here was the two-side 99% confidence interval (CI) of nucleosome-occupied DNA length by MNase-seq data (Cole et al. 2011a; Hu et al. 2014). Last, deNOPA filters candidates for final high-confidence nucleosome positions by a DBSCAN-based outlier detection algorithm (Anant et al. 2010).

**Fig. 1.**
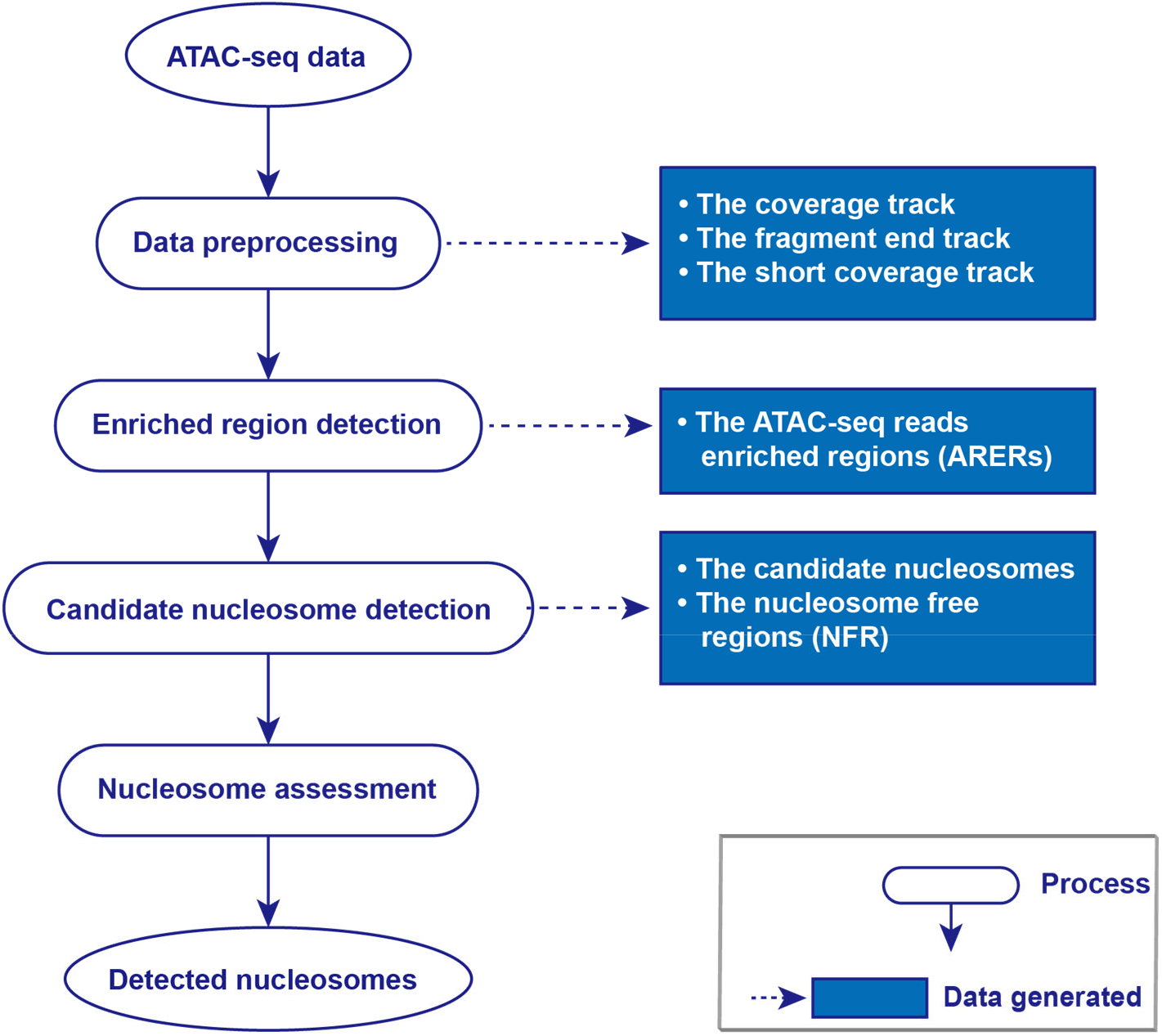
The workflow of deNOPA. Four major steps were summarized in the oval boxes, and the product of each step was indicated by the associated square boxes.

Moreover, deNOPA only predicts nucleosome positions on the so-called ARERs because in most ATAC-seq libraries, about 50%-90% of reads were those mapped in ARERs (Table S1), and the ARERs covered more than about 70% of annotated genes (Table S1), which have most of the predictable nucleosome positions. Details of deNOPA can be found in Methods.

To assess the performance of deNOPA, we compared it with four leading nucleosome positioning detectors, nucleoATAC (Schep et al. 2015), DANPOS (Chen et al. 2013), nucleR (Flores and Orozco 2011) and iNPS (Chen et al. 2014). It is worth noting that nucleoATAC was designed for ATAC-seq in *yeast,* but it can also take pair-end MNase-seq data as input, while DANPOS, nucleR and iNPS were originally designed for MNase-seq data. Following a published definition (Schep et al. 2015), we took the reads with insertion lengths from 101bp to 215bp as the input for these four detectors. We assessed deNOPA with the bulk ATAC-seq data from yeast (Schep et al. 2015), human (Buenrostro et al. 2013) and single-cell ATAC-seq (scATAC-seq) data from mouse (Chen et al. 2018).

### The deNOPA can sensitively and accurately predict nucleosome positions in yeast with ultralow data coverage

We compared the sensitivity, precision and F_1_ scores between deNOPA and other predictors using high-quality, deeply sequenced ATAC-seq data in *yeast* (Schep et al. 2015). The assessment was based on the gold standard chemical mapping data (Brogaard et al. 2012). To assess sensitivity of the predictors, we composed a series of datasets by down-sampling of the raw ATAC-seq reads, which had about 60M read pairs from both long and short fragments, with the sampling rates at 1%, 2.5%, 5%, 7.5%, 10%, 25%, 50% and 75% (Schep et al. 2015).

The sensitivity of deNOPA was consistently high at all sampling rates (Fig. 2A). When the sampling rate was above 50%, i.e., about 30M read pairs, deNOPA, DANPOS, nucleoATAC and iNPS could predict about 60% of chemically mapped nucleosomes, while nucleoATAC quickly became unproductive when the sampling rate decreased. Although DANPOS achieved about 5% higher sensitivity than deNOPA when the sample rates were larger than 1%, deNOPA still outperformed DANPOS at the sparsest condition. At the sparsest condition, the average sensitivity of deNOPA was 90.74% of that with the full dataset, while it was about 70.43% for DANPOS (Fig. 2A). This result implies that the sensitivity of deNOPA is less reliable on reads coverage than DANPOS. We noticed that the nucleosomes specifically mapped by deNOPA were covered by fewer long-fragment reads, implying that short-fragment reads made a considerable contribution towards the greater predictive power of deNOPA at the sparsest condition (Fig. S2A).

**Fig. 2.**
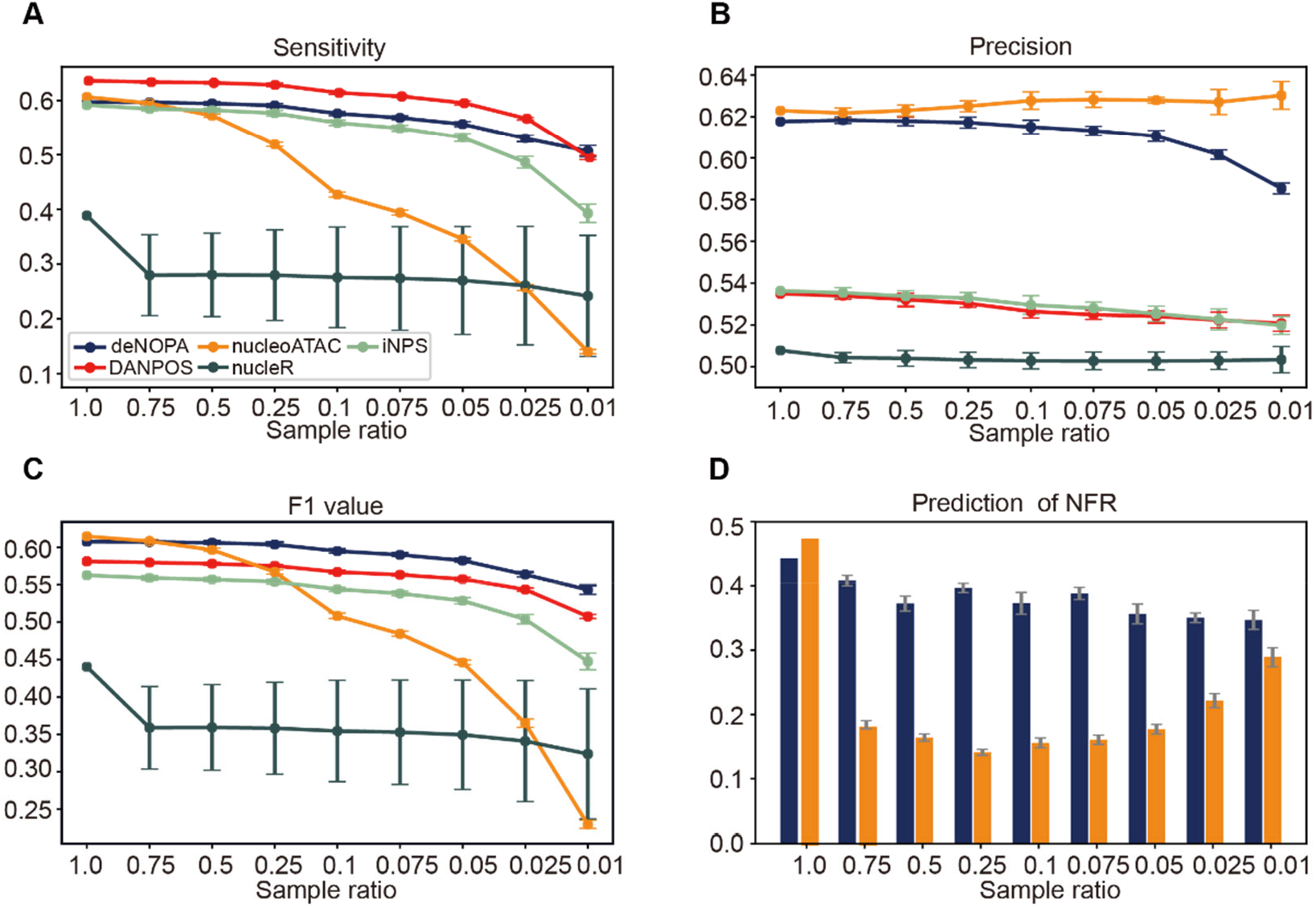
The prediction of deNOPA in yeast was highly accurate. The sensitivity (A), precision (B) and F_1_ value (C) at different sampling rates with data from yeast (Schep et al. 2015). (D) The prediction accuracies of the NFRs of deNOPA (blue) and nucleoATAC (orange). The prediction accuracy was defined as the proportion of predicted NFRs that overlapped with the reference at least one base pair. The error bars were 2 times standard divisions calculated from 10 independent down samples.

The precision of deNOPA was consistently high (Fig. 2B). The precision of a prediction was defined as the relative distance between the predicted and the gold standard nucleosome-occupied region (see SI). We found that the precision of deNOPA and nucleoATAC was consistently higher than the other three MNase-seq tools at all sampling rates (Fig. 2B). It may not be surprising that nucleoATAC had the highest precision, given that it was designed and trained in this dataset (Schep et al. 2015). However, the precision of deNOPA was nearly identical to that of nucleoATAC when the sampling rates were larger than 0.25, and it remained comparable at the lower sampling rates (Fig. 2B). We further assessed precision with MNase-seq data (Cole et al. 2011a) (see SI). Except for nucleoATAC, deNOPA predicted more nucleosome enriched regions with MNase-seq reads compared to all other methods at the lowest sampling rate (Fig. S2B). Using the “nucleosome-to-free ratio” of MNase-seq signals to quantify the enrichment of MNase-seq reads (see SI), we found that it was consistently higher in deNOPA’s prediction than the other tools, even better than that of nucleoATAC (Fig. S2C). Altogether, deNOPA and nucleoATAC demonstrated the highest precision in predicting nucleosome positions in *yeast*.

As a predictor of nucleosome position, deNOPA was able to balance sensitivity and precision. We assessed this balance using the *F_1_* value (see SI). The *F_1_* values of deNOPA were highest when the sampling rate was less than 0.75 (Fig. 2C). The *F_1_* values of nucleoATAC were substantially lower when sampling rates decreased (Fig. 2C).

The deNOPA predicted NFR in the promoter regions better than nucleoATAC at all sampling rates for the *yeast* ATAC-seq data. Since NFR prediction is highly reliant on accurate nucleosome position prediction, better NFR prediction indicates better nucleosome position prediction. Using a public NMase-seq-derived NFR annotation as the reference (Ocampo et al. 2016), we compared the predictions of NFRs between deNOPA and nucleoATAC. We found that the relative accuracy of deNOPA was consistently higher than that of nucleoATAC when the sampling rate was less than one (Fig. 2D). The accuracy of prediction was further evidenced by the enrichment of the DNase-seq signals for deNOPA (Zhong et al. 2016) (Fig. S2D). These results show that deNOPA is a sensitive and accurate nucleosome position predictor with sparse data coverage in *yeast*.

### The deNOPA predicts nucleosome positions well in the human genome

Here, we compared the performance of the predictors with a public ATAC-seq dataset on GM12878 cells (Buenrostro et al. 2013). A total of 153,008 ARERs were detected, utilizing about 57.04% of all sequenced fragments (about 90 million valid read pairs in total) and occupying 32.48% of the human genome. Within those fragments, 47.58% and 52.42% were short and long fragments, respectively. To the best of our knowledge, this was the highest data coverage in the literature at the time of writing this paper. However, it remains an ultrasparse dataset of about 3 × 10^-2^ fragments per base pair, which is equivalent to 0.6% of data coverage in the yeast data (5 fragments per base pair) (Schep et al. 2015). Even after limiting our analysis to ARERs, in terms of nucleosome position prediction, the data coverage remained ultrasparse (about 0.9% compared with that in *yeast*). Thus, we directly assessed the nucleosome callers in ARERs with the full dataset.

The deNOPA was able to predict millions of nucleosomes in human (Fig. 3A). The number of predicted nucleosomes varied substantially from caller to caller. DANPOS, iNPS and deNOPA were the most sensitive tools that predicted 3.01, 3.53, and 1.48 million nucleosomes and covered about 54.95%, 35.24%, and 22.59% of total ARERs in lengths, respectively (Fig. 3A). NucleoATAC and nucleR predicted nucleosomes that covered only 5% and 0.14% of total ARERs in lengths, respectively. Therefore, nucleR was excluded from our further comparison given its insensitivity.

**Fig. 3.**
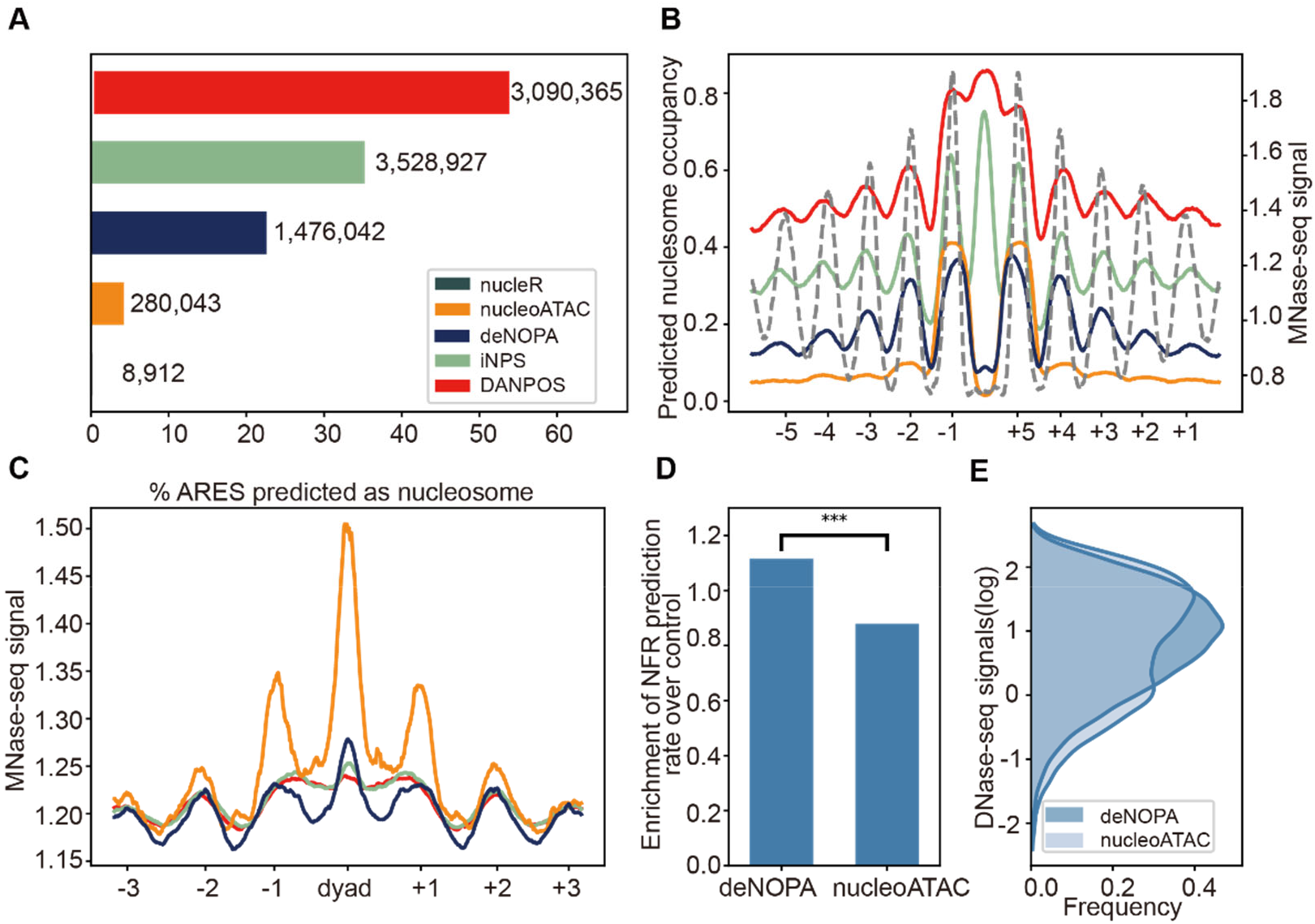
deNOPA predicted nucleosome positions well in human genome. **(A)** The sensitivity of predictors in human GM12878 cells. The x-axis represents the percentage of genome lengths in ARERs predicted as nucleosomes, and the number of predicted nucleosomes is marked by each bar. **(B)** The distributions of predicted nucleosomes around CTCF binding sites. The dashed line represents MNase-seq signals. **(C)** The distributions of MNase-seq signals around predicted nucleosomes. **(D)** The fold enrichment of accurately predicted NFRs. MNase-seq-determined NFRs were taken as standard, and overlapping with at least one nucleotide with the standards was considered a correct prediction. The controls were regions that randomly selected with equal length in the same promoters. **(E)** The distributions of log-transformed DNase-seq signals at deNOPA- and nucleoATAC-predicted NFRs. The orange dashed line marks zero. P-values, *: 10^-2^ ≤ p < 10^-1^ **: 10^-3^ ≤ p < 10^-2^, ***: p < 10^-3^.

Evidence of prediction accuracy in human was stronger for deNOPA than that of the other predictors (Fig. 3B and C). To the best of our knowledge, no chemical mapping data were available for nucleosome positioning in human; therefore, we assessed the precision of the predictions indirectly. First, deNOPA was the only predictor that showed a correct pattern of phased nucleosomes flanking the binding of CTCF and ZNF143 (Fig. 3B and S3A). The prevalence of periodically positioned nucleosomes flanking the binding sites of TFs, e.g., CTCF and ZNF143, is well known (Nie et al. 2014) and has been used to predict chromatin-chromatin interactions (Chepelev et al. 2012). The well-phased nucleosome pattern is commonly seen as flanking TF binding sites in more refined nucleosome position predictions. Based on deNOPA’s prediction, a clear periodic phased nucleosome pattern can be seen flanking the ChIP-seq peak summits of CTCF and ZNF143 in ENCODE (Fig. 3B). However, for DANPOS and iNPS, a nucleosome peak was unexpectedly found at the summit loci of CTCF and ZNF143, (Fig. 3B and Fig. S3A).

Second, compared to deNOPA’s prediction, nucleoATAC’s prediction is biased toward the most well-phased nucleosomes, showing only limited predictive power for the flanking nucleosomes around them. This was first illustrated by its bias toward the two immediate flanking (+1/-1) nucleosomes of the CTCF binding peaks (Fig. 3B). The two nucleosomes flanking CTCF binding peaks consist of 18.50% and 6.74% of the prediction by nucleoATAC and deNOPA, respectively. This was also true for ZNF143, given that 14.92% and 7.51% of predictions flanked the peaks by nucleoATAC and deNOPA, respectively (Fig. S3A). This bias was further evidenced by MNase-seq data (Fig. 3C) (Kundaje et al. 2012). Compared to deNOPA, the contraction of MNase-seq signals in predicted nucleosomes against their neighbors was much higher in nucleoATAC (Pope et al. 2014) (*t* statistics are 28.39 and 48.18, comparing the MNase-seq of predicted nucleosomes and that of their neighbors, respectively, Fig. 3C and S3B). This bias can also be observed in the transcription start site (**TSS**) regions (Radman-Livaja and Rando 2010), i.e., about 3.41% and 0.99% of predicted nucleosomes were +1/-1 nucleosomes around TSSs with nuceloATAC and deNOPA, respectively.

Third, deNOPA predicted nucleosome-occupied regions enriched for active histone modification marks. Histone modifications are chemical groups in the core protein of nucleosomes, while Tn5 preferentially cuts the open and active genome region. Therefore, the active histone modification marks are expected to be enriched in the nucleosome-occupied region (Zhang et al. 2008). Comparing the proportion of predicted nucleosomes that enrich H3K27ac, H3K4me1 and H3K4me3 ChIP-seq signals between the callers (Consortium 2012), we found that the predictions of deNOPA and nucleoATAC had significantly higher enrichment for those histone marks than the other tools and that deNOPA’s prediction was more enriched in H3K27ac and H3K4me1 (Fig. S3C).

Last, deNOPA could accurately predict the NFR in human. Existence of the NFR is a characteristic feature of active promoters, and it can be accurately identified by the pattern of nucleosome positioning (Ocampo et al. 2016). Using MNase-seq data (Kundaje et al. 2012; Ocampo et al. 2016), we defined the NFR in GM12878 cells following the same protocol as that in *yeast* (see SI). Taking these MNase-seq-defined NFRs as a standard, we found more NFRs to be enriched in deNOPA’s prediction than those of nucleoATAC when compared with randomly selected, equal length control regions in the same ARERs (Fig. 3D). The accuracy of NFR prediction was further evidenced by the enrichment of DNase-seq data (Pope et al. 2014) since the NFR enrichment in deNOPA’s prediction was higher than that predicted by nucleoATAC (p = 1.04 × 10^-9^, Mann Whitney U test, Fig. 3E). Furthermore, when comparing the promoters of highly expressed genes (top 20%) with transcriptionally silent (bottom 20%) genes, TSSs showed more overlapping in the deNOPA-predicted NFRs than that of nucleoATAC (the t statistics were 3.91, p = 9.14 × 10^-5^ and 2.41, p = 0.02 for deNOPA and nucleoATAC, respectively, Fig. S3D). Taken together, deNOPA well-balanced the precision and sensitivity in nucleosome position prediction with sparse human ATAC-seq data.

### deNOPA can predict nucleosome positions well at the single-cell level

We assessed the performance of deNOPA at the single-cell level (Chen et al. 2018). As only deNOPA and nucleoATAC showed notable sensitivity and accuracy at ultralow data coverage, we only compared these two tools in this assessment. The testing data were downloaded from the recent work of Chen and colleagues (Chen et al. 2018), which consisted of single-cell ATAC-seq data on five mouse cell types (CD4+ T cell, cardiomyocyte, fibroblast, splenocyte and mESC). To make the data in all cell types comparable, we randomly picked 384 single cells from CD4+ T cell, cardiomyocyte, and splenocyte, respectively, to match the cell number of fibroblast and mESC. To the best of our knowledge, the chemically mapped nucleosomes were only available for mESC (Voong et al. 2016).

The single-cell ATAC-seq data were too sparse to define stable nucleosome positions in individual cells. For a given diploid cell, two copies of DNA, at most, are expected for any loci. About 10.6 million (M) nucleosomes are predicted in the bulk chemical mapping data of mESC cells (Cole et al. 2011a). Thus, at least 20M open linker DNA regions exist in a cell. However, even in a cell with the most abundant reads, only about 0.5M valid read pairs exist per cell (less than 5% of total genome linkers, Fig. S4A). Given the dynamics of nucleosomes, such extremely sparse data provide little help in distinguishing the stable nucleosome positions from the cell-specific transient ones. Therefore, instead of seeking every nucleosome in single cells, we set about assessing how many relatively stable nucleosomes the callers could predict in a pool with a small number of cells, e.g., 384 single cells.

The deNOPA was much more sensitive than nucleoATAC with such pooled single-cell ATAC-seq data in all five cell types. The number of deNOPA-predicted nucleosomes was consistently one or two magnitudes higher than that of nucleoATAC (Fig. 4A). For example, in the smallest dataset, CD4+ T cells, which only contained 9M valid read pairs, deNOPA predicted 145,777 nucleosomes, about three times that predicted by nucleoATAC (45,949). Importantly, the ultrasensitivity of deNOPA was complemented with comparable precision of nucleoATAC, i.e., the precision for nucleoATAC and deNOPA was 0.58 and 0.52, respectively, in mESC, according to the chemical mapping data (Voong et al. 2016) (see SI). The precision and sensitivity of the two algorithms were further assessed by the nucleosome position score (NCP), which indicates the probability that a nucleotide is occupied by nucleosomes (Voong et al. 2016). For algorithms with both high precision and sensitivity, the NCP score should peak at the predicted center position with a periodic pattern for the existence of nucleosome series. Indeed, the NCP peak summits matched well with the predicted nucleosomes by both algorithms, indicating the fine precision for both. However, the periodic pattern of NCP peaks flanking deNOPA’s prediction was much clearer than that flanking nucleoATAC’s prediction, implying the higher sensitivity of deNOPA over that of nucleoATAC in those regions (Fig. 4B).

**Fig. 4.**
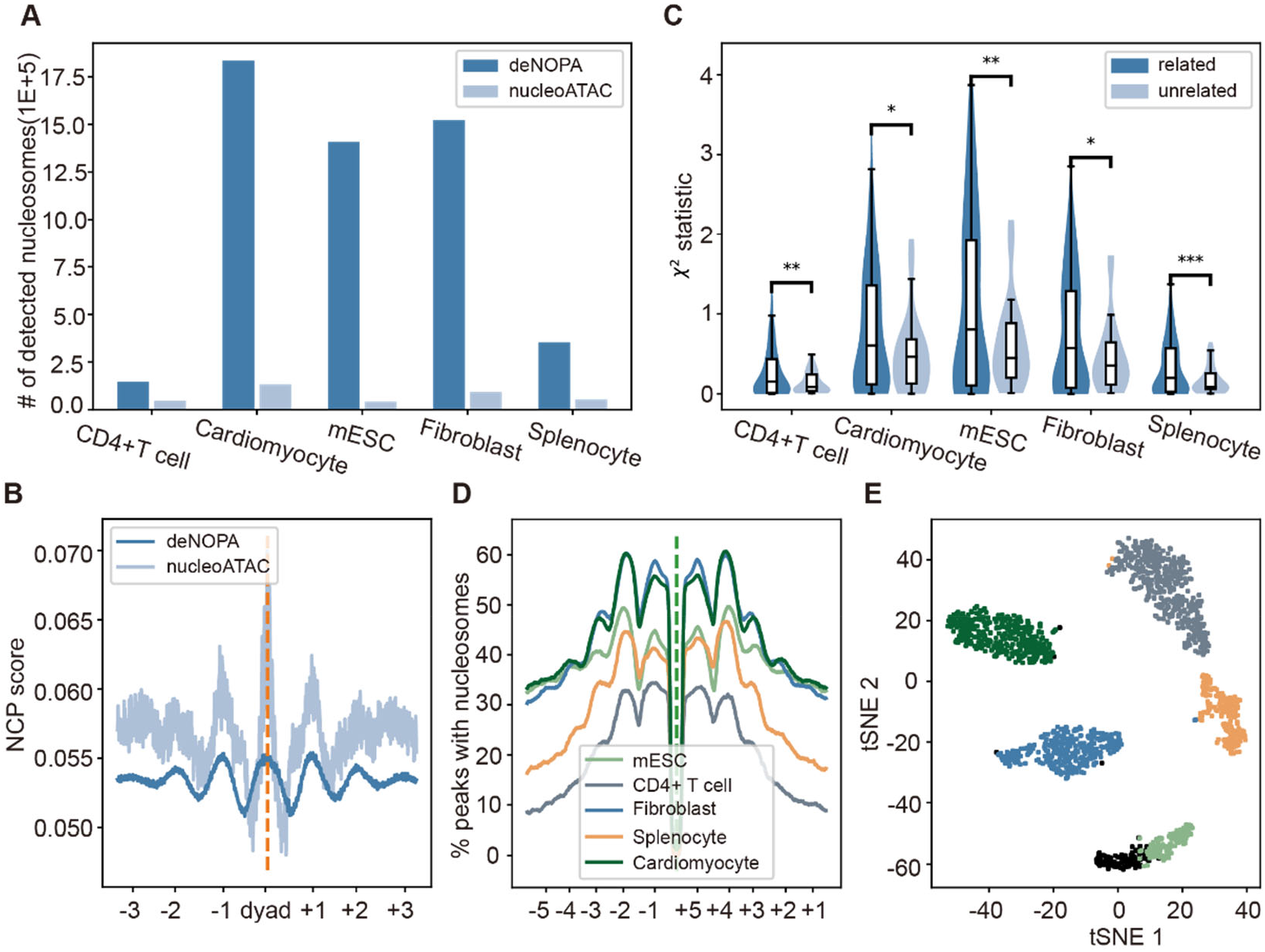
deNOPA-predicted nucleosome positions wellat single-cell level. **(A)** Number of predicted nucleosomes of deNOPA and nucleoATAC in five cell types. **(B)** The distributions of nucleosome center position score (NCP) signals in mESC around the predicted nucleosomes. **(C)** The enrichment of nucleosome-associated 4-mers. The enrichment was indexed by the χ^2^ statistics of comparing the bias of the frequencies of 4-mers at the predicted nucleosome regions and control. The control region was taken from the same ARERs with same lengths of predictions. The differences of the χ^2^ statistic between nucleosome-related and -unrelated 4-mers were tested by Whitney U test. **(D)** The distributions of deNOPA-predicted nucleosome flanking CTCF binding sites. **(E)** The tSNE plot of cell clustering by SCALE based on occupancies of deNOPA-predicted nucleosomes. The misclassified cells were marked as black. P-values, *: 10^-2^ ≤ p < 10^-1^, **: 10^-3^ ≤ p < 10^-2^, ***: p < 10^-3^.

For the other four cell types, we assessed the performance of the predictors indirectly. First, deNOPA’s prediction enriched known characteristic sequence features for nucleosome positioning. For example, higher GC contents were identified from deNOPA’s prediction compared to random controls (Cohanim and Haran 2009; Kaplan et al. 2009) (Fig. S4B). Second, short DNA 4-mers previously reported to bias nucleosome-occupied loci were found to be enriched in deNOPA’s predictions (Tillo and Hughes 2009) (Fig. 4C). This fact implies that deNOPA captures the characteristic sequence of the nucleosome container (Tillo and Hughes 2009). Third, deNOPA-predicted nucleosome positions were periodically arranged flanking the CTCF binding sites. The periodically phased +/-2 and +/-3 nucleosomes could be observed by the stacking of predicted nucleosome positions centered at the CTCF binding sites (Fig. 4D, Fig. S4C and D). The CTCF binding sites were defined by ChIP-seq data and the sequence motif in NFRs for cell types with or without publicly available CTCF ChIP-seq data (Shen et al. 2012), respectively (Fig. 4D and Fig. S4C and S4D). However, only +/-1 nucleosomes flanking the CTCF binding sites were predicted by nucleoATAC (Fig. 4D, S4C and S4D). This indirect evidence supported both the accuracy and sensitivity of deNOPA in nucleosome position prediction.

Collectively, both direct and indirect evidence supported that deNOPA is sufficiently sensitive to predict up to hundreds of thousands of nucleosomes precisely with only a small pool of scATAC-seq data.

### Nucleosome positioning pattern encodes cell identity

It has been shown that accessibility of single cells leads to cell classification (Preissl et al. 2018). Therefore, we asked if deNOPA-predicted nucleosome positioning patterns could also perform this task (see SI). To do this, we took the clusters called from the peaks of MACS2 as reference (Feng et al. 2012) and compared them with the clusters called from the predictions of deNOPA and nucleoATAC. A total of 839137, 363383, and 110964 peaks were predicted in the merged scATAC-seq data of all five cell types by deNOPA, nucleoATAC and MACS2, respectively. By applying SCALE (Xiong et al. 2019), a cell clustering algorithm, to those peaks, we found that deNOPA’s prediction had the highest classification accuracy (Fig. 4E, S4E and F). That is, 84.94% of cells were correctly classified by deNOPA, while 51.94% (p=1.12×10^-96^, one-sided *t* test) and 80.05% (p=2×10^-4^) of cells were correctly classified by MACS2 and nucleoATAC, respectively. We noticed that the classification accuracy of nucleoATAC’s prediction was comparable to that of deNOPA. Given that both deNOPA and nucleoATAC can predict nucleosome positions, our data suggested that nucleosome positions characterize cell identity more clearly than the mixture of nucleosome positions and chromatin accessibilities, as represented by the peaks of MACS2.

Finally, we found that the nucleosome position pattern could be used to identify subcellular lineages from a heterogenous cell population, e.g., tumors. We applied deNOPA and nucleoATAC to a scATAC-seq dataset of human leukemic cell line K562 (Buenrostro et al. 2015) and predicted 1032014 and 68709 nucleosome positions from the 25M uniquely mapped read pairs in 863 cells, respectively. We found that both callers were able to cluster the cells into two groups (Fig. S4G and H). Interestingly, the two groups showed small, but significant, differences between accessibilities, as partially indicated by short fragment reads, on binding sites of key hematopoietic factors, such as GATA1 (*p* = 3.63 × 10^-20^ and 1.89 × 10^-11^ for deNOPA and nucleoATAC, respectively, *t* test) and GATA2 (*p* = 2.03 × 10^-31^ and 6.55 × 10^-56^ for deNOPA and nucleoATAC, respectively, SI, Fig. S4I and S4J). This observation was coincident with two recently reported subtypes of K562 cells characterized as having different accessibilities at the binding sites of those key hematopoietic factors (Litzenburger et al. 2017).

Although the capacity of distinguishing cell subtypes was also observed for nucleoATAC (Fig. S4H), the power was substantially weaker than that of deNOPA (Fig. S4G). Using the inter-to intracluster variance ratio (CH) to index the capacity of clustering, we found that the CHs were 1.06 and 0.58 for deNOPA and nucleoATAC, respectively (Fig. S4G and H), implying a substantially stronger power of clustering for deNOPA. This difference in clustering may stem from the different sensitivities in nucleosome position prediction between the two callers, as 2906 and 38879 nucleosome positions were utilized for clustering in nucleoATAC and deNOPA (see SI), respectively. This example implies that deNOPA may be able to reveal cell population heterogeneity in tumor.

### Application to ultralow coverage of bulk ATAC-seq data of human cells in response to HS

Next, we asked how nucleosome positioning might respond to thermal stress in the context of deNOPA prediction. Using K562 cells as a model system, we performed ATAC-seq experiments under normal (NM, 37°C) and heat (HS, 42 °C, 30 minutes) conditions, and we obtained two high-quality replicate libraries for each condition (SI (Li et al. 2019)). With these data, deNOPA predicted 2,310,244 and 1,280,489 nucleosomes and 114,351 and 90,275 NFRs for NM and HS, respectively, in the ARERs. The accuracy of this prediction was evidenced by the clearly periodic phase pattern of nucleosomes flanking CTCF binding (Fig. S6A and SI). Similar to *Drosophila* (Rowley et al. 2017), the overall accessibilities of NFRs were largely stable after heat shock (Fig. S5 and SI). Thus, we focused on the dynamics of the positions and strength of nucleosomes in response to thermal stress.

First, the position changes of +/-1 nucleosomes flanking the NFR were found to be irrelevant to transcription changes after HS stress. Since +/-1 nucleosomes have been shown to be strongly associated with gene expression regulation (Shivaswamy et al. 2008; Shivaswamy and Iyer 2008; Schep et al. 2015), we might expect changes on either position or strength of those nucleosomes in the promoter of differentially expressed genes (DEGs) after HS. However, little difference in position changes of +/- 1 nucleosomes was found between induced and reduced promoters, not only for their orientation (p = 0.65 and 0.17, for −1 and +1 nucleosome, respectively, *t* test, Fig 5A, Fig. S6C), but also for the distances (p=0.76 and 0.42 for −1 and +1 nucleosome, respectively, KS test). Last, little association was seen between the changes in occupancy of these nucleosomes and changes in the level of expression (p = 0.34 and 0.49 for −1 and +1 nucleosomes, respectively, χ^2^ test, Fig. S6D; see the definition of nucleosome occupancy changes in SI).

**Fig. 5.**
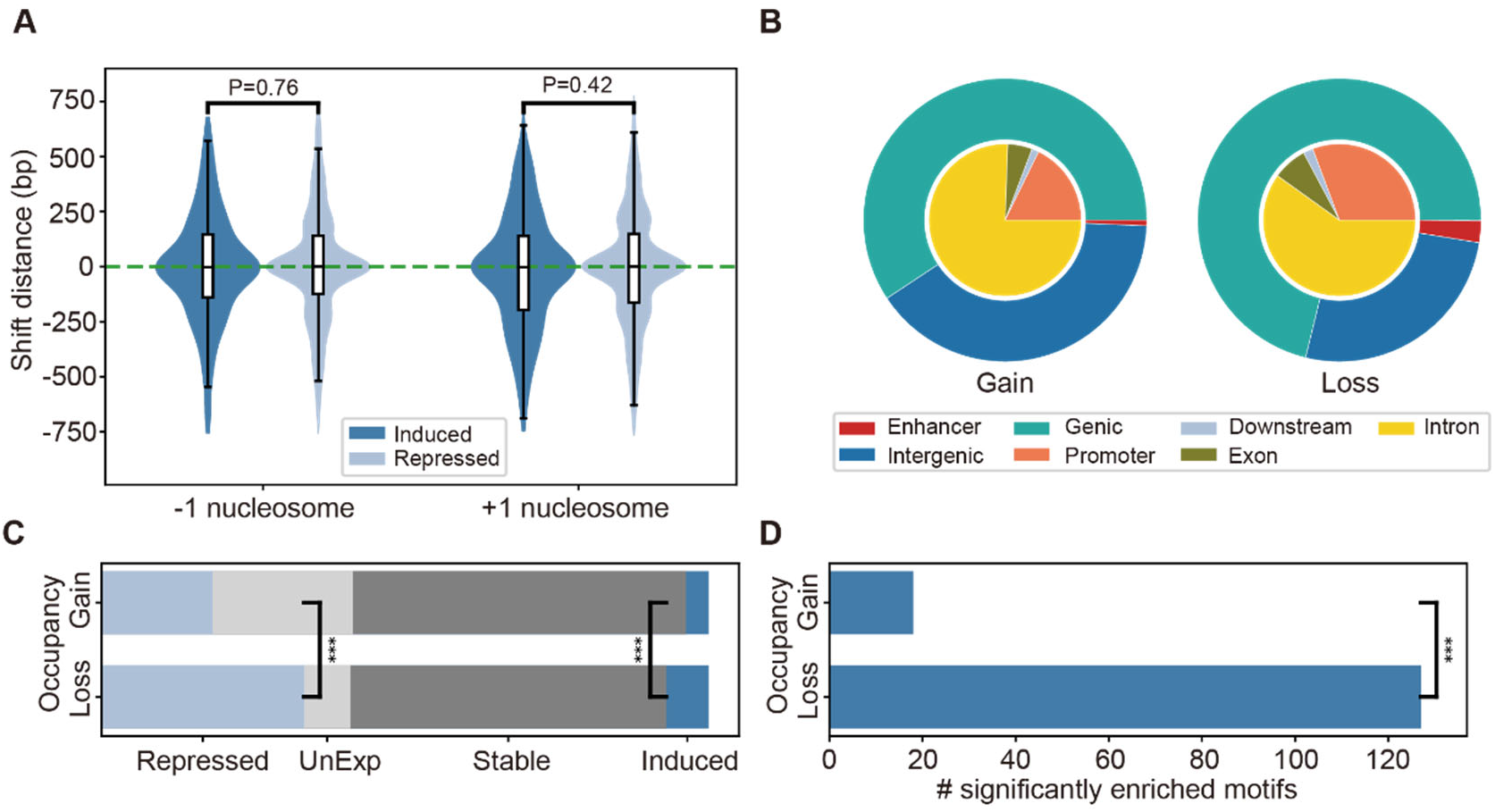
Nucleosome reorganization after K562’s heat shock response. **(A)** The distributions of predicted shifting distance of the +1 and −1 nucleosomes of highly induced repressed genes by deNOPA (KS tests). **(B)** The proportions of nucleosome occupancy of differential regions in regulatory elements. The proportions of nucleosome occupancy to the total number of regions were shown in the outer circle. The proportions of nucleosome occupancy to the number of regions in the intragenic regions were shown in the inner circle. **(C)** The distribution of altered directions of gene expression. The bars gained and lost mark the genes with nucleosome occupancy only gained and lost in promoters, respectively. The genes were grouped according to their change of expression pattern where Repressed, UnExp, Stable, and Induced mark genes with significantly repressed, not expressed, not significantly altered and significantly induced expression level, respectively (*t*-tests). **(D)** The numbers of TF motifs found in the promoter with gained and lost nucleosome occupancy (*t*-tests). ***: P < 10^-3^, **: 10^-3^ ≤ *p* < 10^-2^, *: 10^-2^ ≤ *p* < 10^-1^.

Second, nucleosome occupancy loss was more tightly associated with gene expression changes than nucleosome occupancy gain. Guided by deNOPA-derived nucleosome positions, we detected 68,356 (30.78Mb in total) and 38,751 (15.20Mb) regions that had gain and loss of nucleosome occupancy strength, respectively (see SI). The regions with nucleosome occupancy loss were prone to overlap with intragenic regions or putative enhancers, which were defined by ChIP-seq peaks of H3K27ac located in the intergenic regions (SI). In percentage, 71.15% and 59.31% showed nucleosome occupancy loss and gain in regions that overlapped with the intragenic regions (TSS-2Kb to TTS+3Kb), respectively (p < 10^-52^, *t* test, Fig. 5B). Within the intragenic regions, the regions of occupancy loss were also those most likely to locate at promoters or exons (Fig. 5B). In percentage, 2.56% and 0.56% of nucleosome occupancy was lost and gained in regions that overlapped with the intergenic putative enhancers (Gao and Qian 2019), respectively (p < 10^-52^, *t* test, Fig. 5B). Moreover, the genes with regions of nucleosome occupancy loss solely in promoters (without any region of nucleosome occupancy gained at the same loci) were more likely to be significantly altered after heat shock than those with only regions of nucleosome occupancy gained (Fig. 5C and, Fig. S6E).

Finally, regions with loss of nucleosome occupancy were also those more enriched in transcription factor binding sites than regions with gain of nucleosome occupancy. At the threshold of FDR < 0.05, 127 significantly enriched transcription factor binding motifs occupied regions of loss by HOMER (Heinz et al. 2010), much larger than the 18 in gained regions (p < 10^-4^, *t* test, Fig. 5E and Table S3). Importantly, these factors contained not only key hematopoietic factors, such as GATA1/2 (Lentjes et al. 2016), but also many factors shown to respond to heat shock, such as AP1 and ELF1 (Vilaboa et al. 2017). Such strong biases imply a putative function of nucleosome occupancy loss in response to thermal stress.

When taken together, our data showed that human cells respond to HS with changes in nucleosome occupancy strength.

## Discussion

We developed a sensitive and precise nucleosome prediction algorithm based on ATAC-seq data called deNOPA. Our assessment showed that deNOPA could accurately predict nucleosome positions, even with ultrasparse data that approximated the zone of single-cell ATAC-seq. With hundreds of scATAC-seq data pooled, deNOPA successfully predicted one or two magnitudes more nucleosome positions than existing algorithms. The capacity to predict nucleosome positions with ultrasparse data makes it possible for ordinary laboratories not equipped with sophisticated instruments or reagents to study the dynamics of nucleosomes at the single-cell level. More importantly, with the fast-developing scATAC-seq technologies (Lai et al. 2018; Satpathy et al. 2019), the capacity to predict nucleosome positions with ultrasparse data will also make it possible to study cell-to-cell variations of nucleosome arrangements in the foreseeable future.

The ultrasensitivity of deNOPA mainly stemmed from the additional reads rescued from ATAC-seq short fragments normally discarded in nucleosome-related studies. We found that the cutting sites of short and long fragments are highly correlated, suggesting the rich information carried by short fragments for nucleosome detection. In addition, for the bulk of data utilized by deNOPA, only scantling accuracy is lost to achieve ultrasensitive nucleosome position prediction. It is well known that template searching-based strategies, e.g., nucleoATAC, have high accuracy in pattern recognition (Teif 2016), while model-based methods are generally more sensitive and computationally efficient (Chen et al. 2013; Chen et al. 2014). Although the accuracy of deNOPA was found to be less than that of nucleoATAC in many cases, it is, in fact, comparable to, and much better than, all other competitors we assessed (Fig 2 and 3). Thus, deNOPA can successfully balance sensitivity and accuracy.

It is worth to noting that the current version of deNOPA worked well only with the index-pool-index-based scATAC-seq data, as we showed in this work. However, the distribution of microfluidics-based, e.g., the chromium from 10x Genomics, scATAC data was found to be substantially different from that of index-pool-index-based data in the genome. As clarified in demo data by 10x Genomics, more than 70% of fragments locate at the DHS (Preissl et al. 2018), which is nucleosome-depleted; consequently, few nucleosome positions could be predicted by deNOPA (data not shown).

Finally, we demonstrated the usefulness of deNOPA by applying it on the heat shock response regulation problem of human cells with ordinary ATAC-seq sequencing depth. It has been reported that both chromatin accessibility and nucleosome positions may respond to temperature stress (Shivaswamy et al. 2008; Zeng et al. 2019). However, most of these studies were performed in poikilotherm, in which the temperature in the experiments was relatively lower compared to the mammalian system. One possible reason for the shortage of such research in mammals might involve the temperature sensitivity of micrococcal nuclease (Huang and Garrard 1986). This made deNOPA a good choice to study chromatin and nucleosome dynamics in response to temperature changes in mammals. To the best of our knowledge, the present work is the first such study in the mammalian system on the dynamics of chromatin accessibility and nucleosome positioning in response to heat shock in human cells. The pattern we observed in human cells was similar to that previously reported in fly (Rowley et al. 2017), but different from that of yeast (Lee et al. 2004; Shivaswamy et al. 2008; Shivaswamy and Iyer 2008). A more detailed assay may need to be performed in the future to reveal species-specific regulation of chromatin and nucleosome in response to temperature changes, which trigger more complicated physiological consequences in homothermal animals compared to flies. In summary, deNOPA could be a powerful tool to address broad questions on dynamics of chromatin with ultralow resolution ATAC-seq data.

## Methods

Cell culture and ATAC-seq protocols can be found in SI.

### The deNOPA algorithm

The deNOPA algorithm was mainly divided into four steps: data preprocessing; enriched region detection; candidate nucleosome detection and nucleosome assessment (Fig. 1).

### Data preprocessing

We defined the ATAC-seq coverage of any given genomic locus x as the number of duplicates removed from ATAC-seq fragments covering x, denoted as c(x). The Tn5 cutting events were defined in the +/- 10bp region flanking the 5’ ends of ATAC-seq fragments, and the frequency of cutting events at any given locus x was therefore defined and denoted as s(x). The cutting sites where the raw reads were mapped were shifted towards the center of a fragment by 4-5bp to define the Tn5 recognition sites(Palstra et al. 2003).

The c(x) and s(x) were smoothed by the Gaussian kernel as

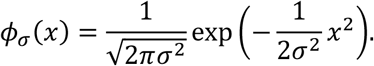

For any given function *y*(*x*), the smoothed function 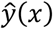 and its derivatives were defined as

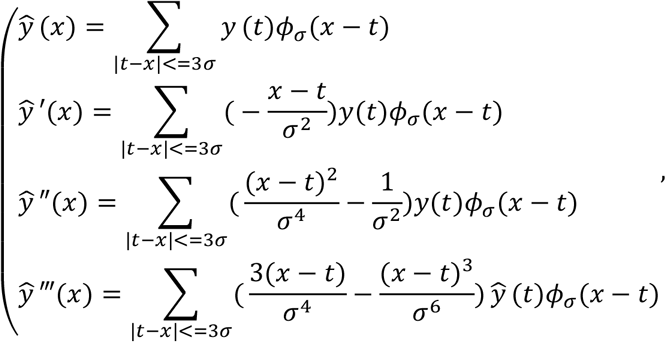

where *σ* = 72 was taken for *c*(*x*), making the smoothing window three times the size of nucleosome to avoid the influence of nucleosome dynamics in the detection of the enriched region, and σ = 24 was taken for s(x), making the smoothing window half the size of the nucleosome to retain enough details of the signal for nucleosome detection.

The distribution of fragment lengths *l* was modeled by a mixture distribution:

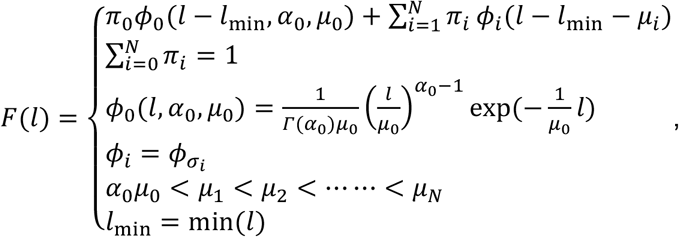

where the fragment length distribution *F*(*l*) was composed of the combination of a Gamma distribution *ϕ*_0_ and a series of Gaussian distributions *ϕ*_1_, *ϕ*_1_,...*ϕ_N_*, where *ϕ*_0_ represented the fragments derived from nucleosome free, accessible regions, and *ϕ*_i_, *i*, = 1,2, …, represented fragment lengths from Tn5 cuts covering *i* nucleosomes. The parameters were estimated by the EM algorithm. The super parameter *N* was determined by maximizing the AIC score. The accessibilities, denoted as r(x), were then calculated in a manner similar to that of c(x), except the fragments were weighted by 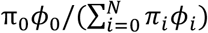, the estimated probability of the fragments generated from the nucleosome free, accessible regions. The r(x) was smoothed with *σ* = 72.

In smoothed 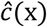 and 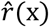, the summits were defined as the genome loci with local maximum. The summits were then modeled by a Gamma distribution, and the parameters were determined by minimizing the mean square error between the modeled 25%, 50% and 75% quantiles and the observations. The significance p-values for summits were assigned according to the modeled distribution. Similarly, the valleys of smoothed 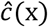 and 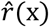, i.e., the local minima, were also modeled by the gamma distributions and assigned p-values accordingly.

### Detection of ATAC-seq reads enriched regions

We only considered the ATAC-seq reads enriched regions (ARERs) in deNOPA, taking the rest of the genome as the background. To detect ARERs, the following steps were applied sequentially:

1. Candidate ARER detection. The ARERs were genome regions with sufficient read coverages in ATAC-seq libraries. To define such regions, small regions with peak read coverages were first determined as candidates. To do this, the summits and their flanking regions with significantly large values of 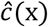 (*p* < 0.1 by default) were termed as the region summits. Genome regions between neighboring region summits without any local minima with small enough value (*p* < 0.5) were then extracted, and those with overlapped bases were merged. Those merged regions were defined as the candidate ARERs.
2. Candidate ARER extension. Each candidate ARER was extended to the nearest local minima in 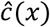 with significantly small value (p < 0.1), both up- and downstream. Neighbor regions were merged if they were near with each other (genome distance < 1kb by default). The final merged regions yielded the ARERs.

### Candidate nucleosome detection

Candidate nucleosomes were detected by detecting the local maximum points in 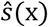.

1. Candidate linker detection: The candidate linkers were roughly defined as those local maximum points of 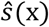 with top 25% of values in the ARERs. The rough candidate linkers were further filtered by the shape of 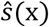 around them. We calculated the genome distance between each rough candidate to its nearest inflection point of 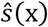 in both directions. The inflection points were defined as 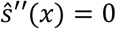 and 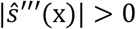. The distances from the local maximum point to the upstream and downstream nearest inflection points were denoted as d_-_ and d_+_, respectively. We show that these distances are between 24-25bp and 15-50bp in the theoretically ideal scenario and 99% empirical CIs from the yeast MNase-seq data, respectively (see SI). Rough linkers with either *d*_-_ or *d*_+_ in the interval between 15 to 50bp were kept.
2. Candidate nucleosome detection: Neighboring rough candidate linkers were grouped to candidate nucleosomes based on the following criteria.

a. The distance between them was similar to the size of nucleosomes. Here 101~215bp was taken as default, which is the 99% CI of the fragment lengths of the yeast MNase-seq data (Cole et al. 2011a; Cole et al. 2011b).
b. The candidate linker upstream should satisfy 15 ≤ d_+_ ≤ 50. At the same time, the candidate linker downstream should satisfy 15 ≤ d_-_ ≤ 50. We define the region between the downstream neighboring inflection point to the upstream linker and the upstream neighboring inflection point to the downstream linker as the inner part of the nucleosome.
3. Determining NFR regions: Regions in ARERs that satisfy the following three criteria were collected as candidate NFRs:

a. Longer than 75bp.
b. No candidate nucleosome overlapped.
c. Overlapped with at least one local maximum point in the smoothed, nucleosome free, accessibility 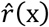. The smallest p-value of the maximum value in these points was taken as the p-value of the NFR (see SI), and the p-values of all NFRs are fed into Benjamini-Hochberg multiple testing adjustment (Benjamini and Hochberg 1995). The NFRs with small adjusted p-value (< 0.1 by default) are reported.

### Nucleosome occupancy assessment

We assess nucleosome occupancy with the following considerations. A stable nucleosome occupancy should cover a length close to 147bp, should be covered by a significantly large number of ATAC-seq fragments, and should have a minimum number of Tn5 cutting events.

The number of ATAC-seq fragments covering the nucleosome was assessed as follows: For a given candidate nucleosome in a specific ARER, the probability of an ATAC-seq fragment covering its whole region, given the condition that the two overlapped, was written as 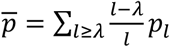, where *p_l_* denoted the proportion of fragments with length *l* in all fragments overlapping the ARER, and *λ* denoted the length of candidate nucleosome. The distribution of the number of fragments, denoted as *θ*, covering the candidate nucleosome was then assumed to follow a binomial distribution 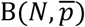, where N denotes total number of fragments overlapping the nucleosome.

The distribution of the number of cutting events located in the inner candidate nucleosome, denoted as *η*, was assumed to follow the Poisson distribution Pois(μτ), where τ denoted the inner length of the nucleosome, and μ denoted the average number of cutting events per base pair in the ARER.

We assessed each ARER separately as a consequence of the large coverage variability of ATAC-Seq data in different ARERs.

Although nucleosome assessment could be conducted based on the p-values calculated from the assumed distributions of segment coverage and the number of cutting events, the assessment could fail under the ultrasparse condition. Alternatively, therefore, we employed an outlier detection-based assessment for ultrasparse data as follows:

The length of candidate nucleosome *λ* was transformed as 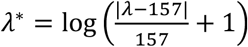, and the number of fragments covering nucleosome θ and the number of Tn5 cutting events in its inner η were transformed by the variance stabilization method (Anscombe 1948; Yu 2009):

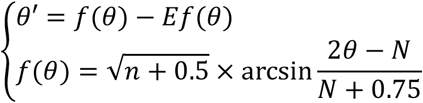

and

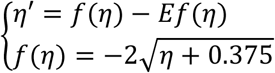

where *Ef*(*θ*) and *Ef*(*η*) were the approximated expectations of *f*(*θ*) and *f*(*η*) calculated by the delta method (Doob 1935). After transformations, *θ’* and *η’* are all distributed with zero means and unit standard deviations, approximately. The larger the transformed value of *θ’* and *η’,* the more probable it is that the nucleosome will be true.

Stable nucleosomes should have large *θ’* and *η’*; therefore, to avoid selecting candidate nucleosomes with large *θ’* or *η’* as outliers, they were then further transformed into *θ\** and *η\** using

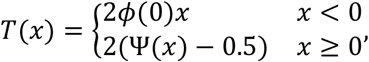

where ϕ and Ψ denoted the probability density function and cumulative distribution function of standard normal distribution. Transformation shrank the distribution at positive half axis towards uniformity, while keeping the shape in the negative side.

A DBSCAN-based outlier detection scheme was then applied among all transformed data (λ*, *θ*\* and *η*\*) using Mahalanobis distances (Anant et al. 2010). The largest cluster was regarded as the predicted nucleosomes.

Two super-parameters are found in the DBSCAN process, the distance specifying the neighborhood, termed eps, and the minimum number of observations in a neighborhood, termed as minpts. Since the distributions of the features differ between different ATAC-seq libraries, these parameters should be calculated specifically for each library. Given two parameters q_1;_ *q*_2_ ∈ (0,1), we calculated eps as the q_1_, quantile of the distribution of minimum distances from each observation to the other ones. Then the number of observations lying in the ball centered at each observation and with radius eps was calculated. The minpts was then calculated as the maximum of these numbers multiplied by q_2_. The default values q_1_, and q_2_ were selected based on the yeast data by optimizing the nucleosome-to-free ratio on the MNase-seq signal profile (see the definition in SI) by grid search. The optimal values *q*_1_ = 0.9 and *q*_2_ = 0.1 were adopted as defaults (Fig. S7).

The outlier detection scheme works because a large part of the candidate nucleosomes consists of true positives. As discussed before, Confidential nucleosomes should have lengths close to the consensus value of 147bp, with a large number of sequenced fragments covered and a few Tn5 cutting events in their inner regions simultaneously, making them naturally closely located to each other in the transformed space and grouped into a large cluster. For those false positives, at least one of the three criteria was overturned, keeping them away from the true positives and acting as outliers to the large cluster composed of true positives.

### Downloaded data sets

For yeast, the ATAC-seq data were download from (Schep et al. 2015) with the GEO accession number GSE63525. Chemical mapped nucleosome positions were downloaded from Table S2 in (Brogaard et al. 2012). The MNase-seq data were downloaded from (Cole et al. 2011a) with GEO accession number GSE26493. The DNase-seq data were downloaded from (Zhong et al. 2016) with GEO accession number GSE69651. For GM12878, the ATAC-seq data were downloaded from (Buenrostro et al. 2013) with GEO accession number GSE47753. Binding positions of CTCF, ZNF143 together with the peaks of H3K27ac, H3K4me1, H3K4me3, the MNase-seq data and the gene expression profile were all downloaded from ENCODE with accession numbers ENCSR000AKB, ENCSR936XTK, ENCSR000AKC, ENCSR000AKF, ENCSR057BWO, ENCSR000CXP and ENCSR000COQ. The DNase-seq data wee downloaded with GEO accession number GSE51334. The mouse single cell ATAC-seq data were downloaded from (Chen et al. 2018) with ENA accession number ERP108537. The chemical mapped nucleosome positions of mESC were downloaded from (Voong et al. 2016) with GEO accession number GSE82127. CTCF binding peaks were downloaded from (Shen et al. 2012) with GEO accession number GSE29184. The single cell K562 data were downloaded from (Buenrostro et al. 2015) with GEO accession number GSE65360. The binding sites of GATA1/2 were downloaded from ENCODE with accession numbers ENCSR000EWM and ENCSR000EWG. Gene expression profiles of K562 cells before and after heat shock were downloaded from (Vihervaara et al. 2017) with GEO accession number GSE89230. Detailed information of these datasets could be found in (Table S1).

## Data access

The ChIP-seq and ATAC-seq data generated in this study have been submitted to the Genome Sequence Archive in the China National Center for Bioinformation (https://bigd.big.ac.cn/gsa/) (Wang et al. 2017) under accession numbers CRA001890 and CRA001891 respectively. The source code for deNOPA can be found at https://gitee.com/bxxu/denopa.

## Acknowledgements

We acknowledge Xiao Li and Ziyang An for useful discussion. Mr. David Martin performed English language editorial services. This work was supported by the Beijing Natural Science Foundation (Z200021), Special Investigation on Science and Technology Basic Resources of MOST, China (2019FY100102), the Beijing Advanced Discipline Fund (115200S001), the Strategic Priority Research Program of the Chinese Academy of Sciences, China (XDA24020307), the National Nature Science Foundation of China (31671342, 31871331, 91940304), and the National Key R&D Program of China (2018YFC2000400).

## Author contributions

XBX and ZZ conceived this project. XBX developed the algorithm. XBX and ZZ analyzed data. XLX, YJ and FFL performed ATAC-seq. XBX, XMG, FFL and ZZ prepared the manuscript. All authors read and approved the final manuscript.

## Supplemental Material

Supplemental_information.docx

Supplemental information and Fig. S1 to S7 for this paper.

Supplemental_Fig_S1.pdf

**Fig. S1.** ATAC-seq reads from short and long fragments have similar information.

Supplemental_Fig_S2.pdf

**Fig. S2.** The accuracy of deNOPA’s prediction in yeast.

Supplemental_Fig_S3.pdf

**Fig. S3.** deNOPA successfully predicted nucleosome positions in human genome.

Supplemental_Fig_S4.pdf

**Fig. S4.** deNOPA successfully predicted nucleosome positions at the single-cell level.

Supplemental_Fig_S5.pdf

**Fig. S5.** Chromatin accessibility remodulation after heat shock of K562 cells.

Supplemental_Fig_S6.pdf

**Fig. S6.** Nucleosome reorganization after K562’s heat shock response.

Supplemental_Fig_S7.pdf

**Fig. S7.** Results of the grid search for parameters **q_1_** and **q_2_** in the DBSCAN process.

Supplemental_Table_S1.xlsx

**Table S1.** Datasets and software packages used.

Supplemental_Table_S2.xlsx

**Table S2.** Cells selected in the scATAC-seq data.

Supplemental_Table_S3.xlsx

**Table S3.** Enriched transcription factor binding motifs in nucleosome occupancy differential regions after K562’s heat shock.

